# B-cell enforced expression of the mouse ortholog of MYD88^L265P^ is responsible for Waldenström-like B-cell lymphoma

**DOI:** 10.1101/794024

**Authors:** Catherine Ouk, Lilian Roland, Alexis Saintamand, Nathalie Gachard, Morgane Thomas, Mélanie Devéza, Nathalie Faumont, Jean Feuillard, Christelle Vincent-Fabert

## Abstract

Here, we created a conditional transgenic mouse model with insertion of a sequence coding for both MYD88^L252P^ and the Yellow Fluorescent Protein (YFP) into the *rosa26*-locus. B-cell specific induction of the transgene constantly led to spleen enlargement with expansion of YFP positive B-cells in 8-12 month-old mice, with a moderate B-cell proliferation increase. Being clonal or oligoclonal, these B-cells exhibited a marked morphological and immunophenotypic lymphoplasmocytic aspect with a plasma cell transcriptomic signature and a serum immunoglobulin M peak. Therefore, continuous activation of MYD88 in mice can lead on its own to a lymphoma close to Waldenström Macroglobulinemia.

**Key point:** B-cell specific enforced expression of MYD88^L252P^ leads to a clonal indolent lymphoplasmocytic B-cell lymphoma with a serum IgM peak.

## Introduction

Accounting for less than five percent of B-cell lymphomas and being an incurable disease, Waldenström macroglobulinemia (WM)/lymphoplasmacytic lymphoma (LPL) is a low-grade B-cell lymphoma primarily located in the bone marrow, with an IgM peak and a lymphoplamaspacytic morphological aspect from the small mature lymphocyte to the terminally differentiated plasma cell^1^. WM/LPLs are genetically characterized by activating alterations of *MYD88*, the most frequent being the *MYD88*^*L265P*^ mutation^2,3^, that leads to canonical NF-κB activation^4^. Consistently, an *in vitro* addiction to MYD88 dependent NF-κB activation has been reported for the BCWM.1 and MWCL-1 WM cell lines^2^.

But Wang et al. showed that retrovirally enforced expression of MYD88^L265P^ in primary mouse B cells is rapidly countered by NF-κB negative feedbacks which may lead to B-cell death^5^. Knittel et al introduced the *Myd88*^*L252P*^ mutation (the murine orthologous mutation of the human *MYD88*^*L265P*^ mutation) into the *Myd88* gene in a B-cell specific manner. These mice occasionally develop spontaneous aggressive B-cell lymphoma when aged, without any IgM peak^6^. Therefore, it is not clear whether continuous activation of MYD88 is responsible on its own of an indolent LPL with an IgM peak or if additional events are required. Here, we present a mouse model in which the MYD88^L252P^ protein was continuously expressed in the B-cell lineage.

## Methods

All material and methods are detailed in supplemental methods.

## Results and Discussion

### The murine MYD88^L252P^ protein constitutively induces NF-κB activation

We first created two transgenes with the mutant murine cDNA sequence of *Myd88* (Myd88^L252P^) or its wild-type form (Myd88^WT^). Both transgenes were in frame with the Yellow Fluorescent Protein (Yfp) sequence, being separated by an Internal Ribosome Entry Site (IRES) sequence (Myd88^L252P^-IRES-Yfp and Myd88^WT^-IRES-Yfp inserts) (Figure 1A). The murine A20 cell line was transfected with a pcDNA3.1 expression vector in which was cloned these two inserts. As shown in Figure 1B and 1C, high levels of MYD88 were produced in both cases and high levels of YFP fluorescence were detected. Cotransfection were performed with the vector containing the luciferase reporter gene downstream either a promoter containing native or mutated NF-κB binding sites (3X-κB-L and 3X-mutκB-L vectors respectively). A strong luciferase activity was detected when MYD88^L252P^ was expressed but not with MYD88^WT^ (Figure 1D). Mutation of the κB binding sites completely abolished the luciferase activity of the transfected cells. This indicates that the MYD88^L252P^ protein was responsible for constitutive NF-κB activation.

**Figure 1:**
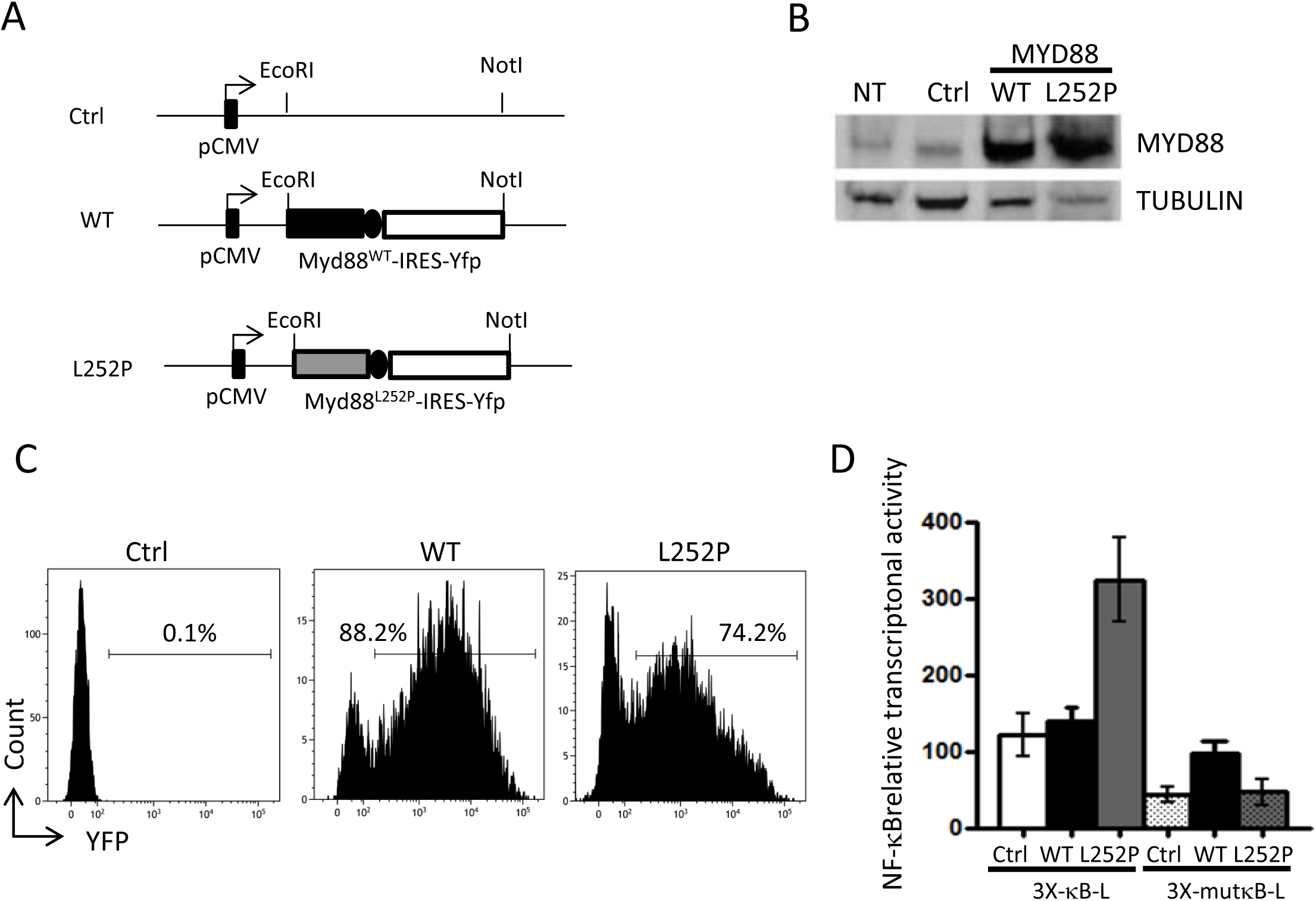
Design of the mouse model. <u>Figure 1A: schematic representation of the transgene:</u> the Myd88^L252P^ sequence was in frame with the Internal Ribosomal Entry Site (IRES) and the coding sequence for Yellow Fluorescent Protein (YFP). This 2.6 kB sequence was directly synthetized and cloned within the pcDNA3.1 vector upstream the pCMV promoter (pcDNA3.1/Myd88^L252P^ vector). As control the same construct with the wild type Myd88 sequence was made (pcDNA3.1/Myd88^WT^ vector). <u>Figure 1 B: MYD88 protein expression after transfection of A20 murine B-cells.</u> Empty pcDNA3.1 (Ctrl), pcDNA3.1/Myd88^L252P^ (L252P) and pcDNA3.1/Myd88^WT^ (WT) vectors were transiently transfected into A20 murine B-cells. Myd88 protein expression of non-transfected (NT) and transfected cells was assessed by Western blot. Revelation of tubulin was used as a loading protein control. <u>Figure 1C: Flow cytometry detection of YFP</u> 48H after transfection of A20 murine B-cells with either the empty pcDNA3.1 (Ctrl), pcDNA3.1/Myd88^L252P^ (L252P) and pcDNA3.1/Myd88^WT^ (WT) vectors. Percentages of positive cells are indicated in each histogram. <u>Figure 1D: Luciferase gene reporter assay:</u> A20 murine B cells were co transfected with either the empty pcDNA3.1 (Ctrl), pcDNA3.1/Myd88^L252P^ (L252P) and pcDNA3.1/Myd88^WT^ (WT) vectors and the Renilla luciferase vector (pRL-TK) which harbors the luciferase reporter gene downstream the CMH class I NF-kappa B binding site (3X-κB-L) or its mutated inactive variant (3X-mutκB-L).

### *MYD88*^*L252P*^ mice develop a splenic clonal indolent lymphoplasmacytic lymphoma with an IgM peak

The Myd88^L252P^-IRES-Yfp insert was subcloned downstream the neomycin-STOP cassette flanked by LoxP sites of the pROSA26.1 vector^7^ (Figure 2A). After transfection of ES cells and neomycin selection, cells with the transgene were injected in C57BL/6J blastocyst. Stable germinal transmission of the Myd88^L252P^-IRES-Yfp transgene was obtained (MYD88^L252P-*flSTOP*^ mice). Spleen B cells of these mice were *ex vivo* treated with the TAT Cre recombinase. Expression of YFP was detected in the vast majority of the cells while no background was found in absence of TAT Cre (Figure 2B).

**Figure 2:**
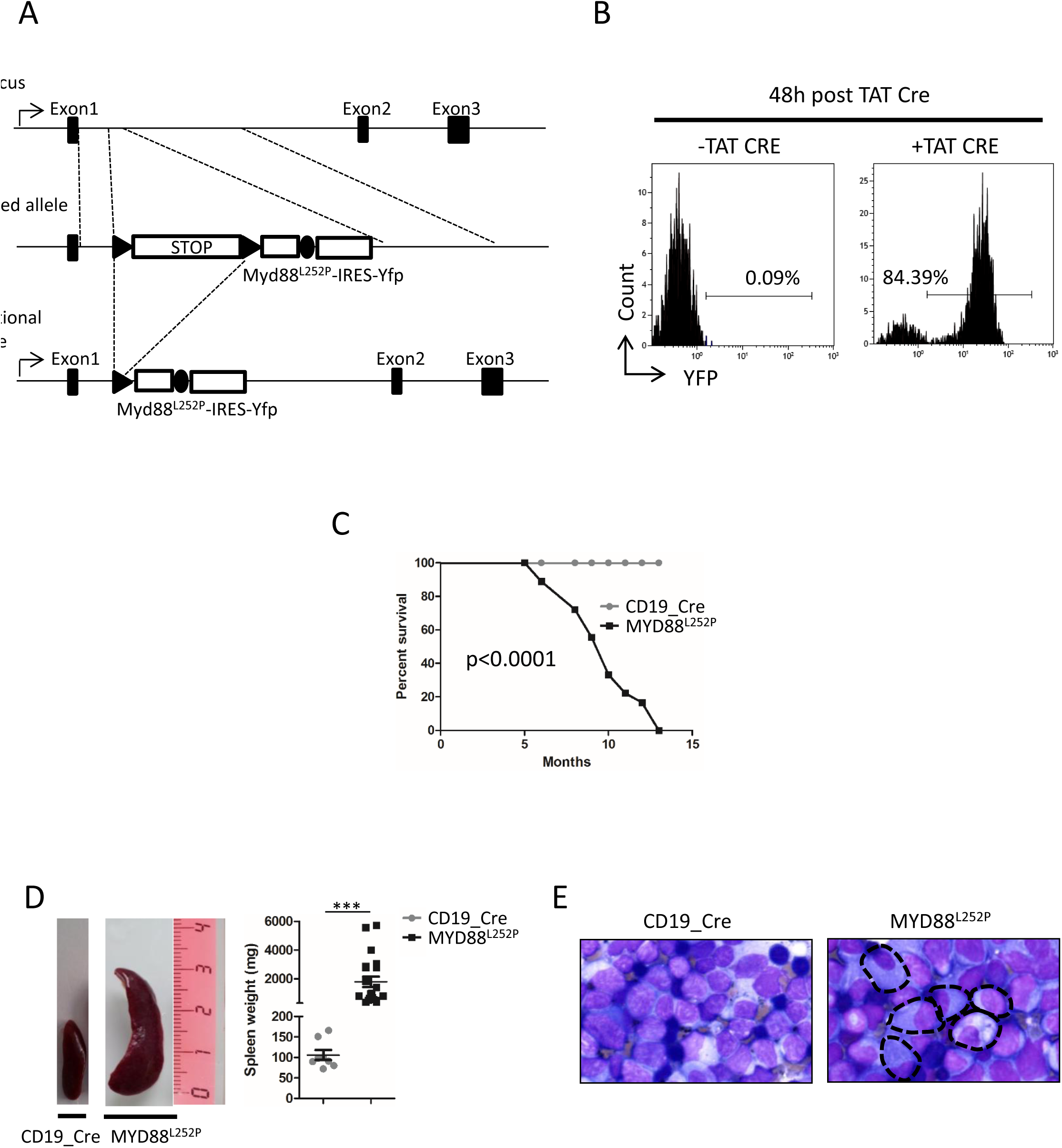
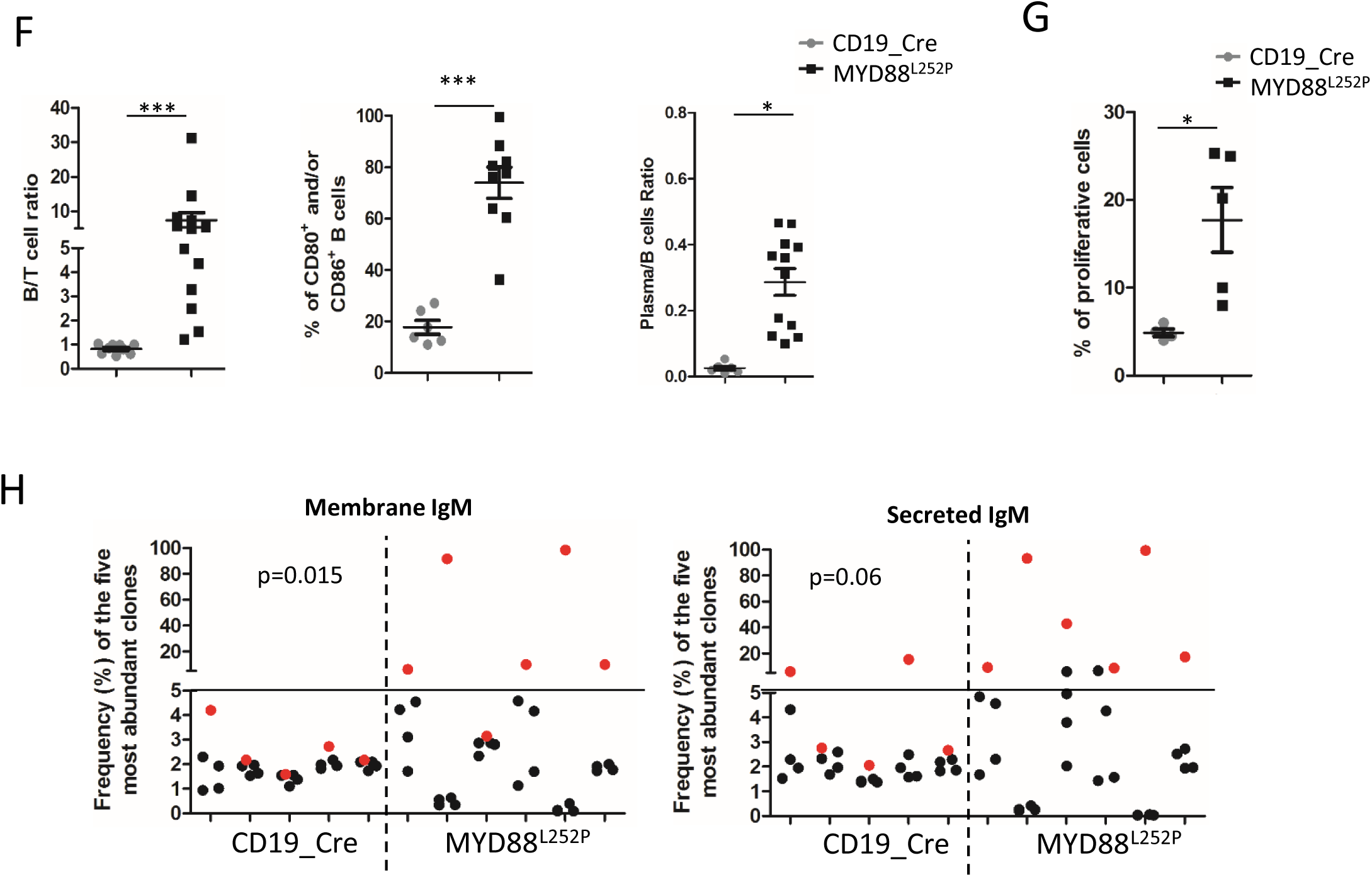
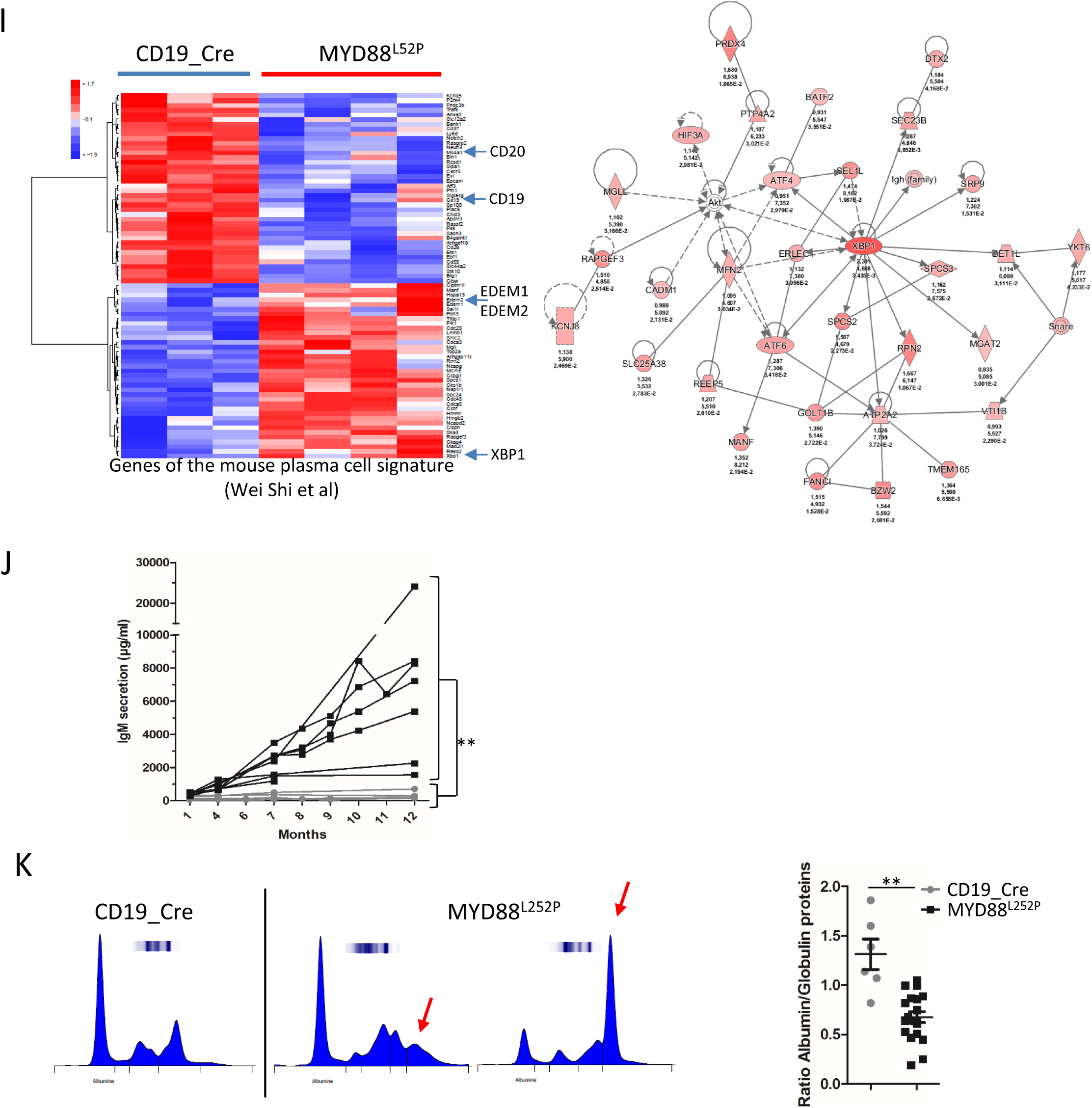
MYD88^L252P^ transgenic mice developed lymphoplasmacytic B cell lymphoma with engagement in plasma cell differentiation and secretion of a IgM peak. <u>Figure 2A: Targeting strategy for the insertion of a conditional MYD88 L252P-IRES-YFP (MYD88^L252P-*flSTOP*^) into the murine *rosa26*-locus.</u> This strategy was similar to the one published by Hömig-Hölzel et al^12^. Briefly, Cre mediated recombination leads to deletion of the stop cassette and expression of MYD88 L252P-IRES-YFP (MYD88^L252P^ mice) under transcriptional control of the endogenous *rosa26*-promoter. STOP: Stop cassette. <u>Figure 2B: Flow cytometry detection of YFP in purified B cells from spleen of MYD88^L252P-*flSTOP*^ after in vitro induction of the Cre recombinase.</u>Purified B splenocytes were incubated 2 hours incubation with or without the TAT Cre recombinase, washed twice in culture medium and cultured for 48 hours. <u>Figure 2C: Kaplan Meyer survival curve of CD19_Cre and MYD88^L252P^ mice. MYD88^L252P^</u> mice were sacrificed at the terminal stage of their disease. CD19_Cre littermates were used as control. <u>Figure 2D: Spleen size of CD19_Cre and MYD88^L252P^ age-paired mice.</u> MYD88^L252P^ were sacrificed at the terminal stage of their disease together with at least one CD19_Cre mouse of the same age. Left panel: example of the spleen macroscopy; right panel: histogram of the spleen weights (mean and standard deviation are indicated for each mouse group; ***: p<0.001). <u>Figure 2E: Cell morphology of CD19_Cre (upper panel) and MYD88^L252P^ (lower panel) splenocytes.</u> Spleen imprints were May-Grünwald stained according to routine hematological procedures. Cells with a marked plasma cell differentiation are dot line surrounded cells (see also Supplemental Figure 1). <u>Figure 2F: histograms of spleen B/T cell ratio (left panel), percentages of activated spleen B cells (middle panel) and YFP+ plasma cells/YFP^+^ B cells ratio (right panel) from CD19_Cre and MYD88^L252P^ mice</u> (*: p<0.05; ***: p<0.001). <u>Figure 2G: In vivo proliferation index of spleen B cells from CD19_Cre and MYD88^L252P^ mice.</u> BrdU was intra peritoneally injected 18 hours before sacrifice. Percentage of BrdU positive B cells was assessed by flow cytometry (*: p<0.05). <u>Figure 2H: µ heavy chain B-cell clonal abundance analysis:</u> clonal abundance was evaluated from high-throughput sequencing of VDJ regions after mRNA reverse transcription and a RACE PCR with primers specific for either the membrane (left panel) or the secreted (right panel) form of the mouse μ heavy chain. The relative frequency of the five more abundant clones is shown for five and six CD19_Cre and MYD88^L252P^ mice respectively (left and right part of the graph respectively). The most abundant clone is highlighted in red. <u>Figure 2I: gene expression analysis of the plasma cell signature:</u> total mRNA was extracted from whole spleen tissues. Gene expression profiles were obtained using the MoGene-2_1-st-v1 Affymetrix chip. One thousand five hundred fifteen mRNA transcripts were selected to be differentially expressed between CD19_Cre and MYD88^L252P^ mice and homogeneously expressed within each mouse type. Among these genes, 77 belonged to the plasma cell signature published by Wei Shi et al^11^. Left panel: hierarchical clustering of genes belonging to the plasma cell signature. Right panel: deduced XBP1 regulatory network using the Ingenuity Pathway Analysis software for genes upregulated in MYD88^L252P^ mice belonging to the “<u>13 22 4 23 39 7 24</u>” aggregated Kmean cluster (see Supplemental Figure 4 and supplemental methods for detailed explanations). <u>Figure 2J: IgM serum levels of CD19_Cre and MYD88^L252P^ mice:</u> kinetics of IgM serum levels in CD19_Cre and MYD88^L252P^ mice (**: p <0.01). <u>Figure 2K: serum protein electrophoresis (SPE) for CD19_Cre and MYD88^L252P^ mice.</u> Left panel: typical SPE examples for one CD19_Cre (left) and two MYD88^L252P^ (middle and right) mice. Monoclonal peaks are pointed by the red arrow. Right panel: histogram of the serum albumin/globulin ratio of CD19_Cre and MYD88^L252P^ mice (**: p<0.01).

MYD88^L252P-*flSTOP*^ and CD19_Cre mice^8^ were crossed together. Mice with both transgenes (MYD88^L252P^ mice) had a median life expectancy of 9 month, with a continuously decreasing overall survival (Figure 2C). At nine to twelve month-old, MYD88^L252P^ mice had a marked splenomegaly with increased spleen weight (Figure 2D). Morphologically, cells showed a lymphoplasmacytic aspect (Figure 2E) with massive infiltration of the spleen white pulp by a mixture of small lymphocytes, lymphoplasmacytic and plasma cells (Supplemental Figure 1). By flow cytometry, B/T-cell ratio was increased in all MYD88^L252P^ mice (Figure 2F, left panel). This increase was associated with constant up-regulation of CD80 and/or CD86 activation markers (Figure 2F, middle panel and Supplemental Figure 2A). Partial engagement in plasma cell differentiation was immunophenotypically evidenced by expression of the CD138 plasma cell marker on a significant proportion, but not all, YFP positive B-cells (Figure 2F, right panel) with concomitant decrease of B220 expression (Supplemental Figure 2B). Proliferation index was constantly increased (Figure 2G) albeit being always below 30%. MYD88^L252P^ tumors had the same µ heavy chain mRNA clonal expression of both secreted and membrane form, as evidenced by high-throughput sequencing of VDJ regions after mRNA reverse transcription (Figure 2H). As control, no significant B-cell clonal expression of the mouse γ heavy chain was found (Supplemental Figure 3).

With a fold change of 2, a set of 1515 differentially expressed genes were found in MYD88^L252P^ spleen tumors when compared to spleen of CD19_Cre mice (Supplemental Table 1). This included seventy seven deregulated genes that belong to the plasma cell signature published by Shi *et al*^9^ (Supplemental Table 2). Specific unsupervised clustering with these 77 genes revealed that MYD88^L252P^ tumors exhibited features of plasma cell differentiation such as decreased expression of *Cd19* and *Msa4* (*Cd20*) or increased expression of *Xbp1* (Figure 2I left panel). By combination of K-mean and hierarchical clustering with principal component analysis, deregulated genes in MYD88^L252P^ spleen tumors were segmented in 14 classes of co-regulated genes (see supplemental methods, Supplemental Figure 4A and Supplemental Tables 2 and 3 for details). Functional analysis of these classes showed that genes belonging to the T-cell lineage, as well as T-cell signaling and activation signatures were down-regulated in MYD88^L252P^ spleen tumors (Supplemental Figure 4B). At the opposite, expression of genes belonging to the proliferation, *RelA Nf-kB* and plasma cell differentiation signatures were up-regulated (Supplemental Figure 4B). Indeed, a network of 34 up-regulated genes centered on *Xbp1* could be revealed (Figure 2I, right panel).

In the blood, IgM levels continuously increased from month 4 to 12 (Figure 2J) and an IgM peak was systematically found in serum of 8 – 12 month old MYD88^L252P^ mice, with constant decrease of the albumin/globulin ratio (Figure 2K).

Beside the fact that primary B-cells were *ex vivo* retrovirally infected before reinjection, the model of Wang et al^5^ raises the question whether the human MYD88^L265P^ protein may have exactly the same activation properties in mouse than in human B-lymphocytes. *Myd88* endogenous regulation is poorly known. But *Myd88* methylation status is important in glioblastoma^10^ and an alternative splice variant of *Myd88* unable to activate NF-κB has been reported^11^. The model of Knittel *et al*^6^ raises the question of the relationship between B-cell lymphomagenesis and regulation of the endogenous *Myd88* locus throughout B-cell life. By inserting our transgene in the *rosa-26* locus, we enforced continuous expression of the mutated Myd88 transgene in a heterozygous-like context while respecting the native mouse MYD88 activation pathways. Thereby, we created a novel conditional MYD88^L252P^ mouse model which constantly developed a splenic indolent B-cell lymphoma with lymphoplasmacytic differentiation and secretion of a monoclonal IgM. This resumes three of the four main characteristics of WM. Primary localization to the bone marrow was not observed. Whether this relies in human and mouse niches differences or in the lack of additional genetic events such as CXCR4 mutation remains to be determined. Nevertheless, our results genetically demonstrate that continuous activation of MYD88 is able to promote clonal emergence of a B-cell lymphoma very closed to WM. This provides an interesting preclinical model for development of new therapeutic approaches in MW or to study the immune surveillance for example. Indeed, a better understanding of the underlying molecular mechanisms of WM is necessary to in turn be exploited in order to develop new specific therapeutics specifics for B-cell cancer.

## Acknowledgements

The team of J. Feuillard is supported by grants from the Ligue Nationale Contre le Cancer (Equipe labellisée Ligue), the Comité Orientation Recherche Cancer (CORC), the Nouvelle Aquitaine Region and the Haute-Vienne and Corrèze committees of the Ligue Nationale Contre le Cancer. We thank the animal facility of the University of Limoges for their assistance. We are indebted to the laboratory of immunology of the Hospital University Centre of Limoges for mouse serum protein electrophoresis. We are extremely grateful to Marion Espeli and Karl Balabanian (team “Chemokine regulated interplays between lymphocytes and their environment” Inserm U1160 “EMiLy” Institut de Recherche Saint-Louis, Paris, France) for in depth scientific discussion of our results. We also thank Olivier Bernard (INSERM U1170, Institut Gustave Roussy, Villejuif, France) for scientific exchanges and advices.

## Authorship Contributions

C.O. and L.R. contributed equally to this work. C.O. and L.R. performed and analyzed experiments. A.S. helped to perform the repertoire analysis. M.D. an N.G. performed the transcriptomic experiments. J.F. A.S. and L.R. made the bioinformatics analyses. N.F. participated to the design of the project. C.V.F. created the mouse model, contributed to the experiments and analyzed the results. J.F and C.V.F. wrote the manuscript. All authors read and approved the final manuscript.

