## Supplemental Figure for "B-cell enforced expression of the mouse ortholog of MYD88^L265P^ is responsible for Waldenström-like B-cell lymphoma"

### Supplemental Figure 1

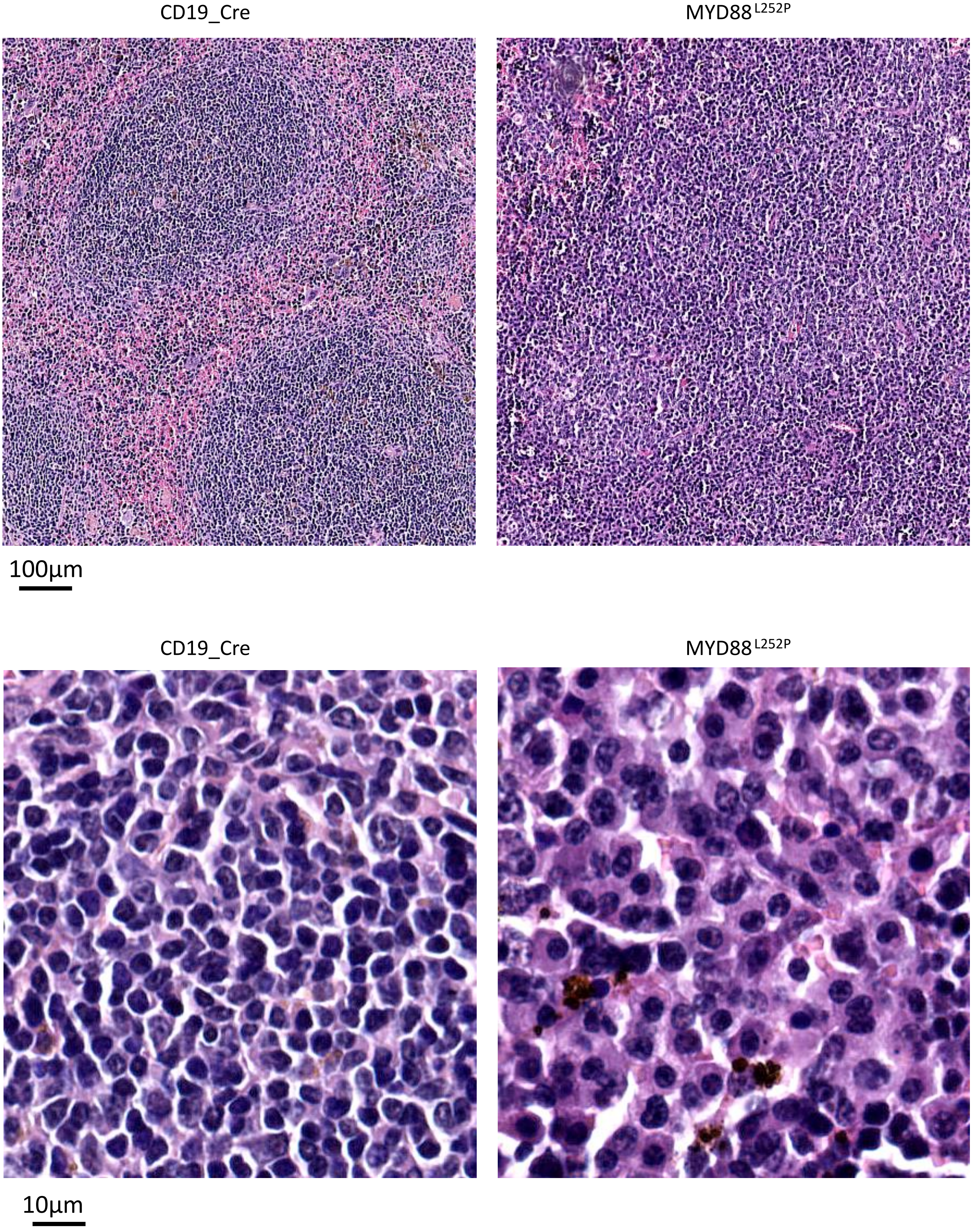

### Supplemental Figure 2A

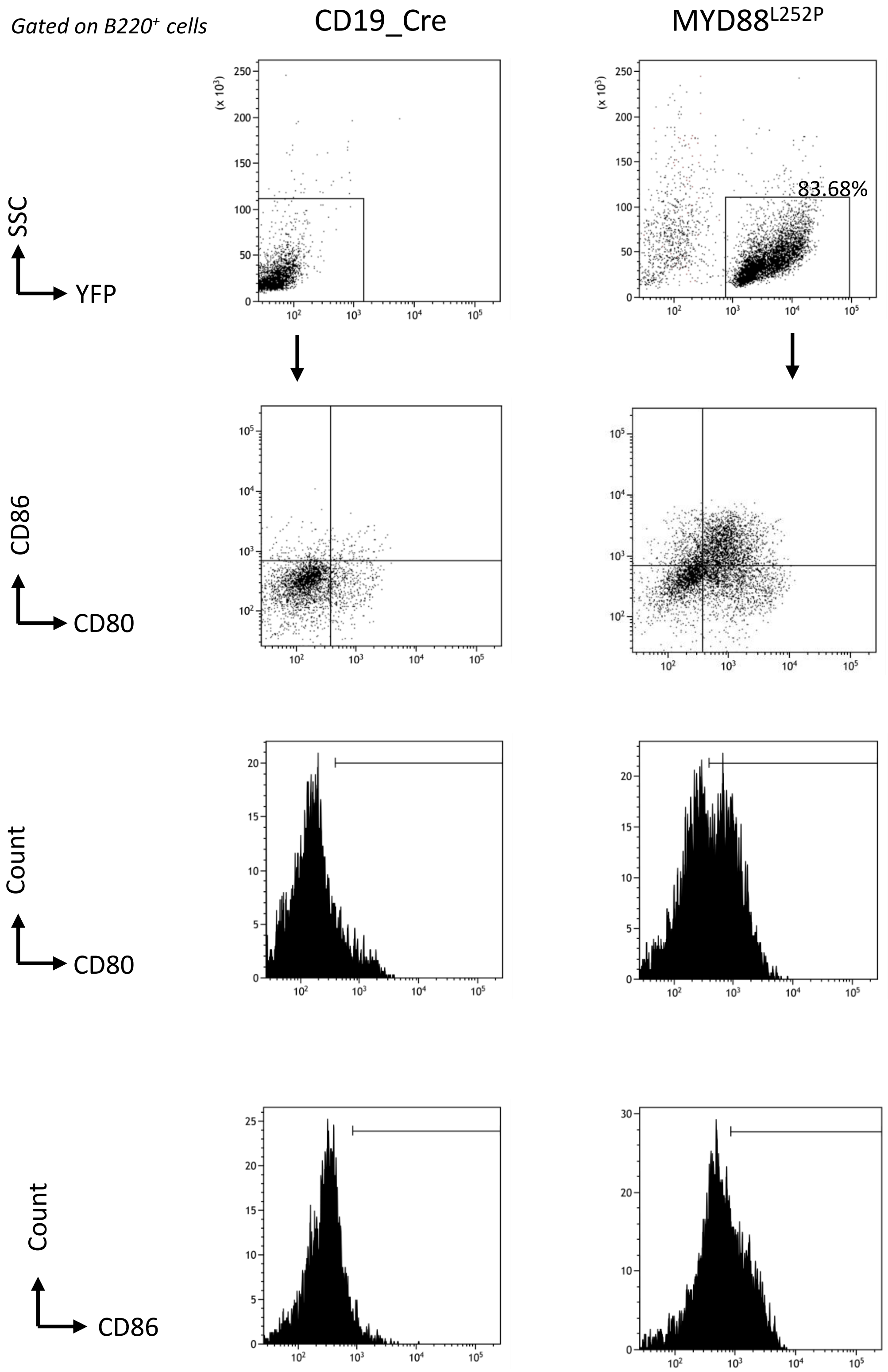

### Supplemental  Figure 2B

*Gated on CD3⁻ cells*

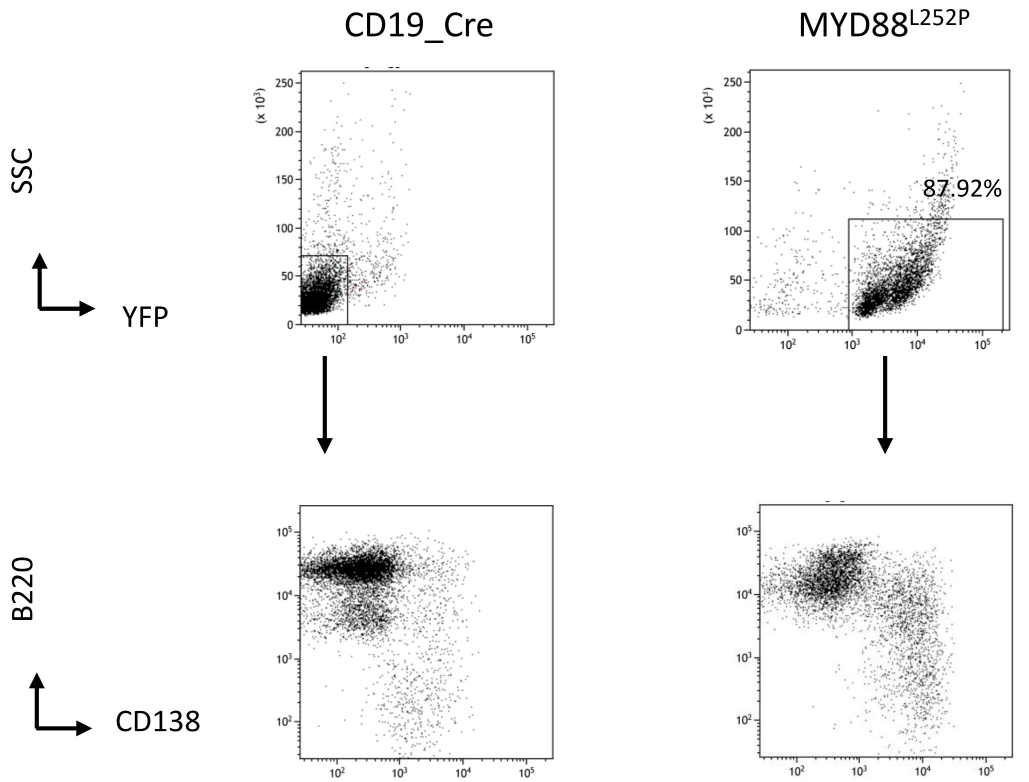

Supplemental Figure 3

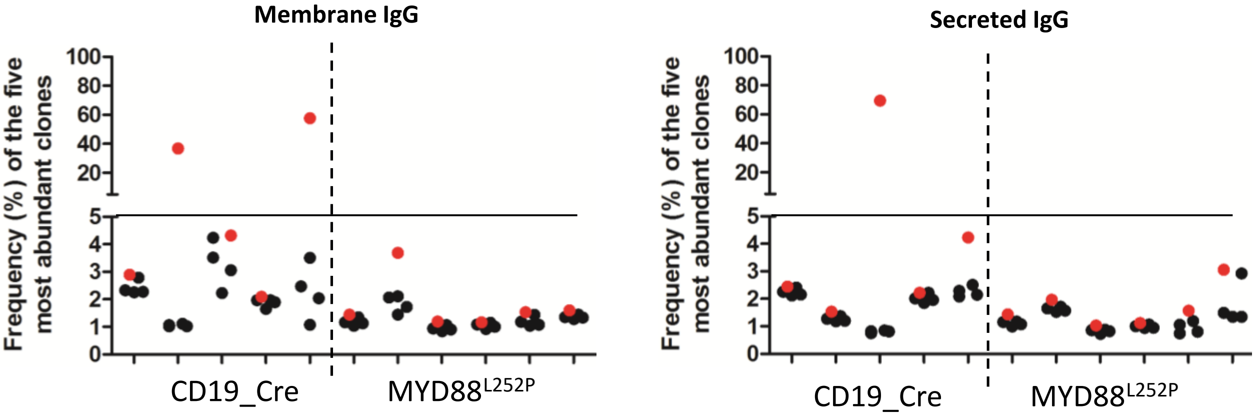

### Supplemental Figure 4A

1515 gènes
40 clusters

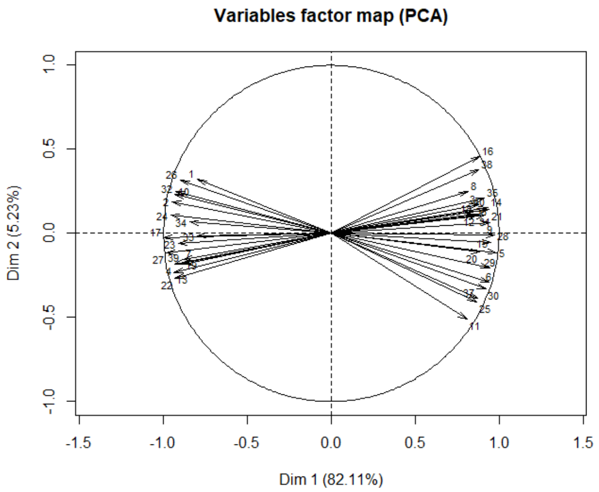


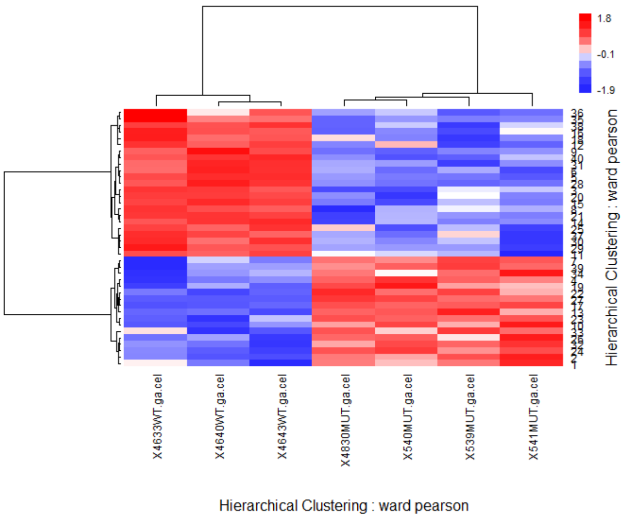


1515 gènes
14 clusters

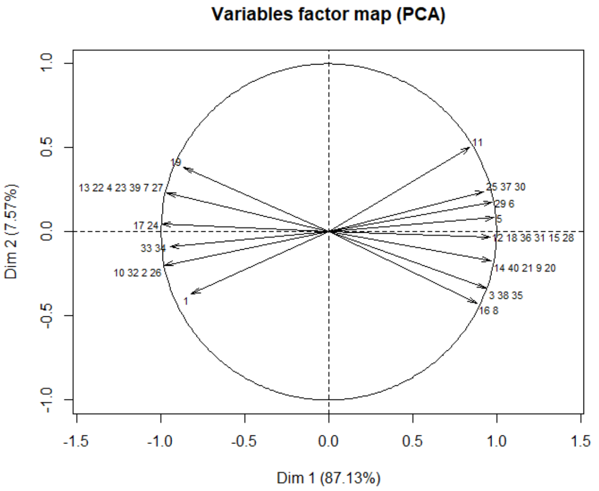


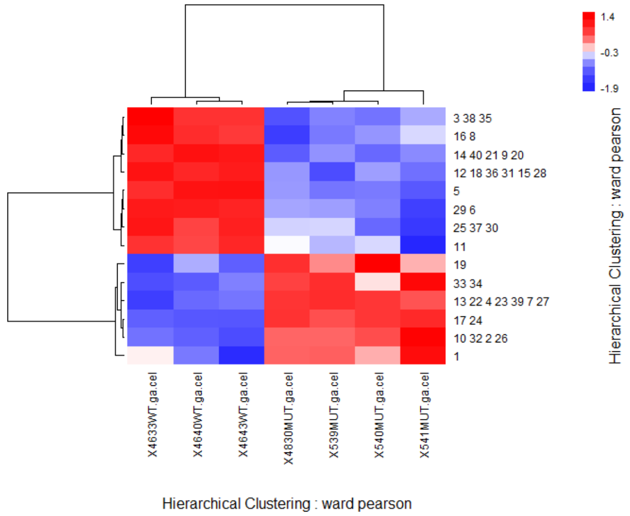

### Supplemental Figure 4B

| Aggregated KmeanCluster | Pathway/function |
|---|---|
| **5** n=41 | SP1↘ |
| **33 34** n= 54 | Myc↗ Oct2↗ TP53↘ |
| **3 38 35** n=94 | STAT1↘ SPIB↘ |
| **29 6** n=79 | CEBP A and B ↘ STAT4 ↘ T-cell activation ↘ |
| **25 37 30** n=86 | TP53↘ RELB↘IL4↘ |
| **19** n=11 | XBP1 ↗ |
| **17 24** n=154 | Myc↗ E2F1 and 3↗ TP53↘ Cell proliferation ↗ Proteasome ↗ |
| **16 8** n= 65 | TGFB1↘ IL1B ↘ |
| **14 40 21 9 20** n=128 | EBF1↘ RELB↘ EZH2↘ T-cell signature and activation↘ |
| **13 22 4 23 39 7 24** n= 314 | E2F1↗ RELA↗ XBP1↗, MYD88↗ Proliferation↗ Plasma cell diff↗ Metabolism↗ |
| **12 18 36 31 15 28** n=187 | IL4↘ BCL6↘ T-cell signaling↘ |
| **11** n=36 | NFATC2↘ IL18↘ TH1 and TH2 activation ↘ |
| **10 32 2 26** n=184 | E2F1↗ Myc↗ Cell Cycle↗ |
| **1** n=28 | MAZ↗ Proteine ubiquitination ↗ |

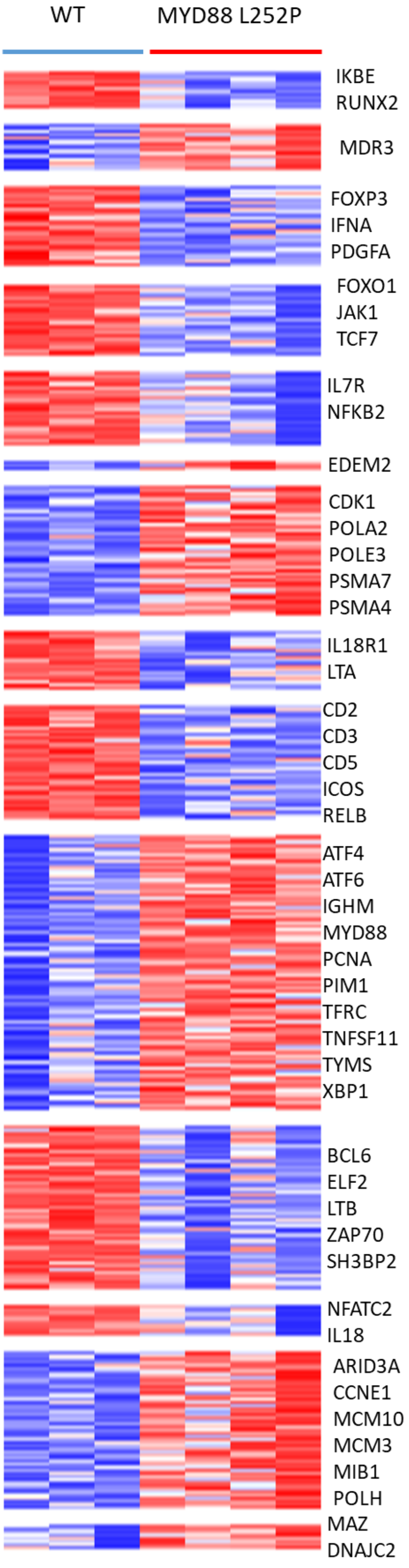
